# Nucleic Acid Aptamers Protect Against Lead (Pb(II)) Toxicity

**DOI:** 10.1101/2024.03.28.587288

**Authors:** Afreen Anwar, Solimar Ramis De Ayreflor Reyes, Aijaz Ahmad John, Erik Breiling, Abigail M. O’Connor, Stephanie Reis, Jae-Hyuck Shim, Ali Asghar Shah, Jagan Srinivasan, Natalie G. Farny

## Abstract

Lead (Pb(II)) is a pervasive heavy metal toxin with many well-established negative effects on human health. Lead toxicity arises from cumulative, repeated environmental exposures. Thus, prophylactic strategies to protect against the bioaccumulation of lead could reduce lead-associated human pathologies. Here we show that DNA and RNA aptamers protect *C. elegans* from toxic phenotypes caused by lead. Reproductive toxicity, as measured by brood size assays, is prevented by co-feeding of animals with DNA or RNA aptamers. Similarly, lead-induced behavioral anomalies are also normalized by aptamer feeding. Further, cultured human HEK293 and primary murine osteoblasts are protected from lead toxicity by transfection with DNA aptamers. The osteogenic development, which is decreased by lead exposure, is maintained by prior transfection of lead-binding DNA aptamers. Aptamers may be an effective strategy for the protection of human health in the face of increasing environmental toxicants.

**SYNOPSIS:** Lead remains a pervasive environmental contaminant with significant human health implications. This study investigates an entirely novel intervention for the problem of lead toxicity, using nucleic acid aptamers.

## INTRODUCTION

Major water crises like the notorious case of Flint, MI^1^, and the ongoing problem of lead (Pb(II)) contamination in school water systems across the U.S.^2^ make clear that lead exposure remains a pervasive public health problem. Lead causes neurodevelopmental anomalies in children^3^ and increased risk of cardiovascular disease (CVD)^4^, renal damage, and neurological disease in adults^5,6^. Lead exposures are cumulative, and the neurologic damage caused is permanent. The molecular mechanism of lead toxicity is related to its ability to replace calcium in biological processes ^7^. Lead enters cells through calcium channels and can interfere with calcium ion flow, which is a central mechanism of its neurotoxic effects^8^. Lead integrates into hydroxyapatite and can be stored for many years in bone^9^, leaching back into the body even after exposures are eliminated^10,11^ and thus taking years or decades to be fully cleared from the body. Deposition of lead in the bone also interferes with bone cell signaling, maturation, and differentiation^12–14^, leading to impaired fracture healing^15^ and increased risk of osteoporosis^16,17^.

Even very low levels of lead have been associated with health risks. Blood lead levels as low as 6.7 μg/dL in adults were associated with higher all-cause mortality, CVD mortality, and ischemic heart disease mortality^18^; even levels as low as 2 µg/dL represent a significant CVD risk^19^. Blood lead levels in children less than 5 µg/dL are associated with neurodevelopmental problems such as decreased academic achievement, decreased IQ, and increased incidence of attention-related and problem behaviors^20^. These findings suggest that the Centers for Disease Control and Prevention (CDC) guidelines for actionable blood lead levels, currently 3.5 µg/dL for children^21^ and 5 µg/dL for adults^22^, are insufficient to protect against the negative health consequences of lead exposure. While intravenous EDTA chelation therapy is being explored to lower CVD risk in high-risk populations including diabetics^23^, there is not currently an approved, cost-effective, minimally invasive prophylactic or therapeutic intervention for chronic low-dose lead poisoning, which affects millions of people worldwide^24^.

Aptamers are short stretches of nucleic acids (<100 nucleotide single-stranded DNA or RNA molecules) that bind specifically to a target molecule or ion^25^. There are numerous aptamers currently in clinical trials for diseases ranging from various cancers to autoimmune conditions^26^ and one aptamer drug (Macugen) is approved for the treatment of macular degeneration. Thus, as a therapeutic modality, aptamers have great promise^27^. Several studies have identified lead-binding aptamers with high sensitivity and specificity using Systematic Evolution of Ligands by Exponential Enrichment (SELEX)^28^. One such study identified an aptamer known as Pb7S with low micromolar affinity (1.60 ± 0.16 µM) for lead ions and reported minimal cross-reactivity to other ions^29^. Structural analyses suggest that lead may associate with nucleic acids through G-quadruplex (G4) structures^30^, though the mechanism of lead ion interaction for Pb7S has not been determined. Most lead-binding aptamers, including Pb7S, have been developed for the purpose of lead detection in environmental and biological samples^28^. Whether lead-binding aptamers could be used clinically as specific lead ion chelators for the prophylactic or therapeutic treatment of lead poisoning has not yet been reported.

The nematode *Caenorhabditis elegans* is a ∼1 mm transparent soil organism that has been used as a laboratory model organism for decades. The well-established genetic tools, facile imaging, rapid life cycle (∼2 weeks), ease of maintenance, and complex-yet-mapped neural network^31,32^ make *C. elegans* an ideal organism for a broad array of topics from aging^33^ to neurodegeneration^34^ to behavior^35^. These features have made *C. elegans* an ideal model organism for toxicology studies as well. Meta analyses have revealed that *C. elegans* toxicology studies have been just as predictive of adverse human outcomes and mechanisms of toxicity as rodent studies^36^, and yet can be performed at a fraction of the expense in time and resources.

Here we show, using a *C. elegans* model, that lead-chelating DNA and RNA aptamers applied in the presence of lead protect the animals from reproductive and behavioral toxicity. Both DNA and RNA versions of the aptamers are effective, and the protective effect is specific to lead and to the aptamer sequence. Further, using both HEK293 and primary murine osteoblast culture models, we show that aptamers protect cultured cells from lead toxicity, and protect osteoblastic function. The results suggest aptamer-based chelation could be further developed as a prophylactic or therapeutic strategy for human exposures to toxic metals.

## RESULTS

### Lead Ions Interact Specifically with Lead-Binding ssDNA Aptamers

The lead-binding single stranded (ss)DNA aptamer Pb7S (Table 1 and Supplementary Table S1) has been described previously^29^. This aptamer was originally selected for use in a fluorescence-based detection assay for lead (Supplementary Fig. S1)^29^. The aptamers were 5’ end labeled with a fluor (fluorescein amidite, FAM), and annealed to a shorter antisense strand with a 3’ quencher (dabcyl, DAB), which quenched the fluorescent output. Upon aptamer binding to lead ions, the quench strand was released, resulting in a fluorescent signal. To confirm lead binding to the Pb7S aptamer, we reproduced the fluorescence-based lead binding assay, testing combinations of two flours (FAM and Yakima Yellow) and three quenchers (DAB, Black Hole Quencher 1 (BHQ1), and Iowa Black (IAB)) (Supplementary Fig. S2). Using the Yakima Yellow fluor, we found a statistically significant difference in fluorescence from the no lead control at 20 µM lead, indicating an interaction of lead with the aptamer. The fluorescence detection system was highly sensitive to pH, with lower (pH 5.5) and higher (pH 8.4) values resulting in a loss of dynamic range (Supplementary Fig. S3), which we suspected was a caveat of using fluorescent detection, rather than a pH-dependent association of lead with the aptamer.

**Table 1:**
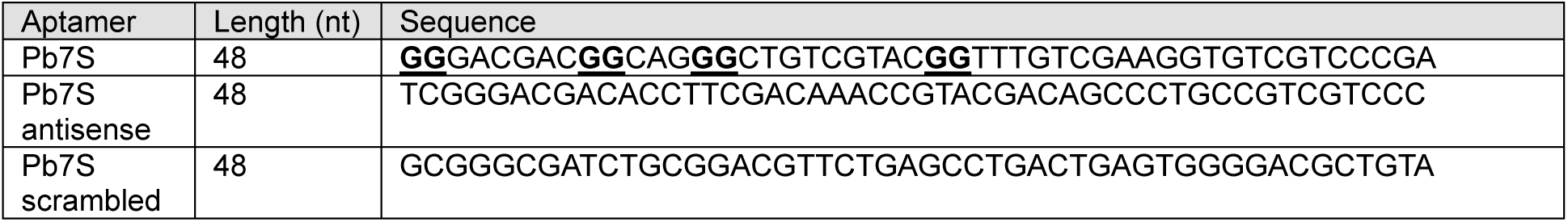
Sequences of Pb7 aptamers and controls. Putative G-quadruplex residues shown in bold and underlined.

Structural modeling predicts the formation of a G-quadruplex (G4) structure in the Pb7S aptamer^37^ (Table 1). Lead ions are known to assemble into G4s with high affinity^38^, creating unique G4 signatures by circular dichroism (CD) spectroscopy^39^. To confirm the specific interaction of the Pb7S aptamer with lead ions in a manner independent from fluorescence detection, we applied CD to measure the lead-dependent assembly of the G4 (Fig. 1). The unbound Pb7S aptamer, and antisense and scrambled controls, displayed a strong CD maximum in a single peak at approximately 285 nm. The addition of lead ions to the Pb7S aptamer resulted in the concentration-dependent appearance of a broad peak with a maximum at 317 nm (Fig. 1A).

**Figure 1.**
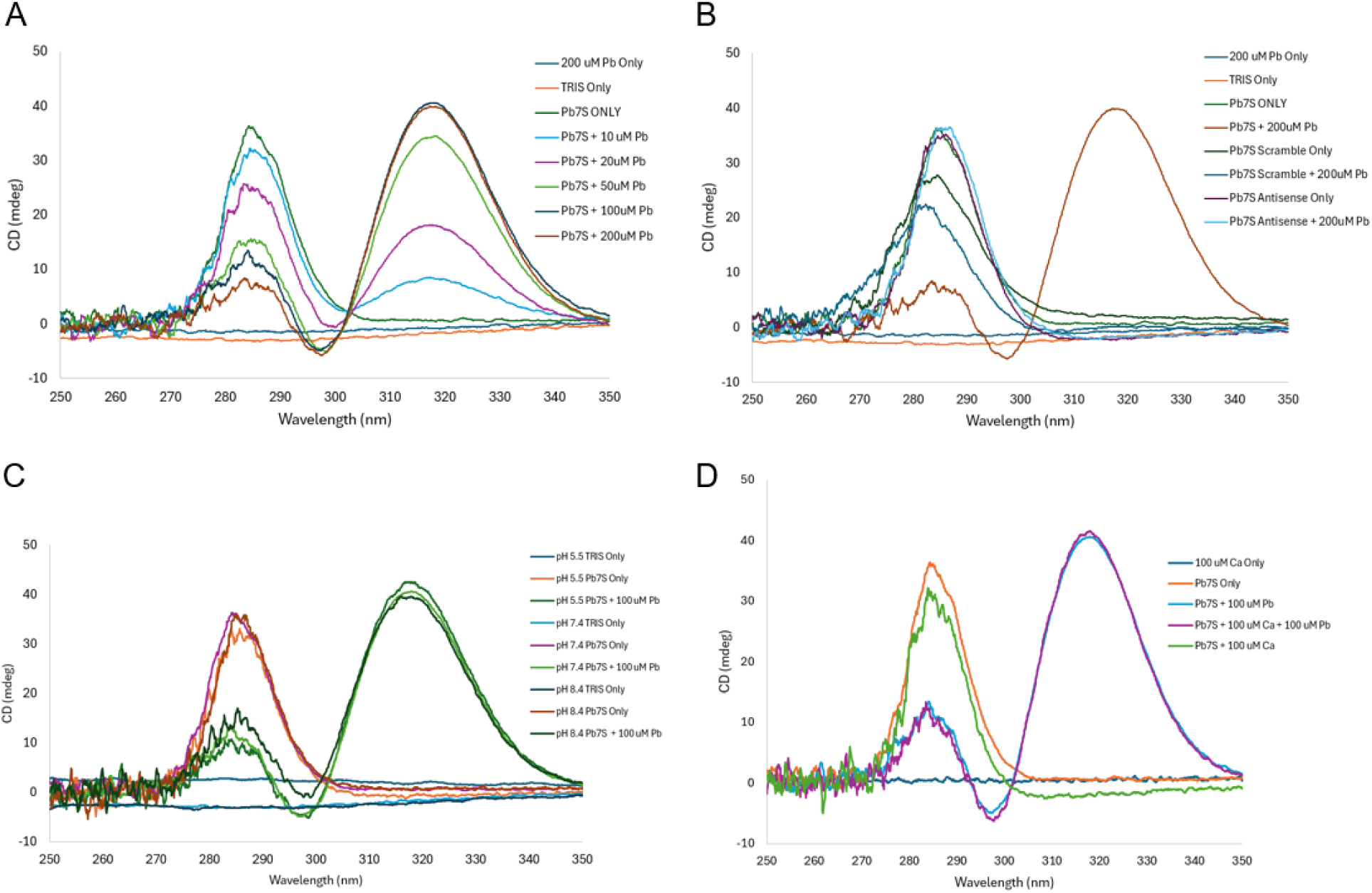
Pb7S binds specifically to lead ions. A) Lead acetate (Pb) was added at indicated concentrations to 10 µM samples of DNA aptamers (Pb7S) in 10 mM Tris pH 7.4. B) Pb7S antisense or scrambled aptamer controls (10 µM) were mixed in 10 mM Tris pH 7.4 with or without 200 µM Pb as indicated. The same graph of the Pb7S aptamer with 200 µM Pb from panel A is overlaid in panel B for reference. C) 10 µM samples of Pb7S aptamer, with or without Pb as indicated, were mixed in 10 mM Tris pH 5.5, 7.4, or 8.4. D) 10 µM samples of Pb7S aptamer, with or without 100 µM Pb and/or 100 µM calcium, as indicated, were mixed in 10 mM Tris pH 7.4. For all samples, CD measurement was performed on a Jasco J-1500, scanning at 100 nm/minute, 20 accumulations/sample, from 250 nm – 350 nm in 0.1 nm intervals.

Antisense aptamer and scrambled aptamer controls did not form the 317 nm peak in the presence of lead (Fig. 1B). The formation of the peak at 317 nm was identical when the pH of the solution used was 5.5 or 8.4, suggesting that the interaction is not particularly sensitive to pH in this range (Fig. 1C). As lead mimics calcium within biological systems, we investigated the binding of the Pb7S aptamer to calcium, and found no evidence of binding of the Pb7S aptamer to calcium (Fig. 1D). Further, the presence of 100 µM calcium did not alter the formation of the 317 nm peak when lead was added subsequently. From our CD experiments, we conclude that the Pb7S aptamer binds with high specificity to lead ions, not calcium.

### Reproductive Toxicity of Lead is Prevented by Pb-Binding ssDNA Aptamers

Lead has previously been shown to result in reproductive toxicity in *C. elegans*^40–42^, causing a dose-dependent decrease in brood size. These prior studies were conducted with animals exposed to metals by continuous growth in liquid cultures in multi-well plates. To better mimic dietary exposure to metals, we exposed our animals to metals by feeding. We first confirmed the dose-dependent decrease in brood size using our experimental feeding method. L3 stage animals were plated to NGM agar seeded with their food source OP50 *E. coli* mixed with lead acetate at concentrations from 0 – 25 mM. We found by this method that 15 mM lead exposure mixed in the OP50 lawn was sufficient to reliably cause a minimum 50% decrease in brood size (Supplementary Fig. S4).

To determine whether chelation of lead ions with aptamers could reduce reproductive toxicity, we employed three strategies to expose the animals to the aptamer: feeding, soaking, and drop casting (Supplementary Fig. S5A, S6A, Fig. 2A). The feeding strategy mixed the aptamers at the designated concentration into the OP50, with or without lead, then animals were plated to this mixture and offspring were counted (Supplementary Fig. S5A). The soaking strategy exposed animals to aptamers in an aqueous solution for 2.5 hours, then the animals were moved to NGM plates seeded with OP50 with or without lead (Supplementary Fig. S6A). For the drop casting method, animals were plated to NGM plates containing OP50 with or without lead, then 10 µL of aptamer at the indicated concentration was dropped onto the animal (Fig. 2A). In all cases, the *C. elegans* cuticle is not permeable to the aptamer, and thus ingestion of the aptamer is the predicted mechanism of exposure.

**Figure 2.**
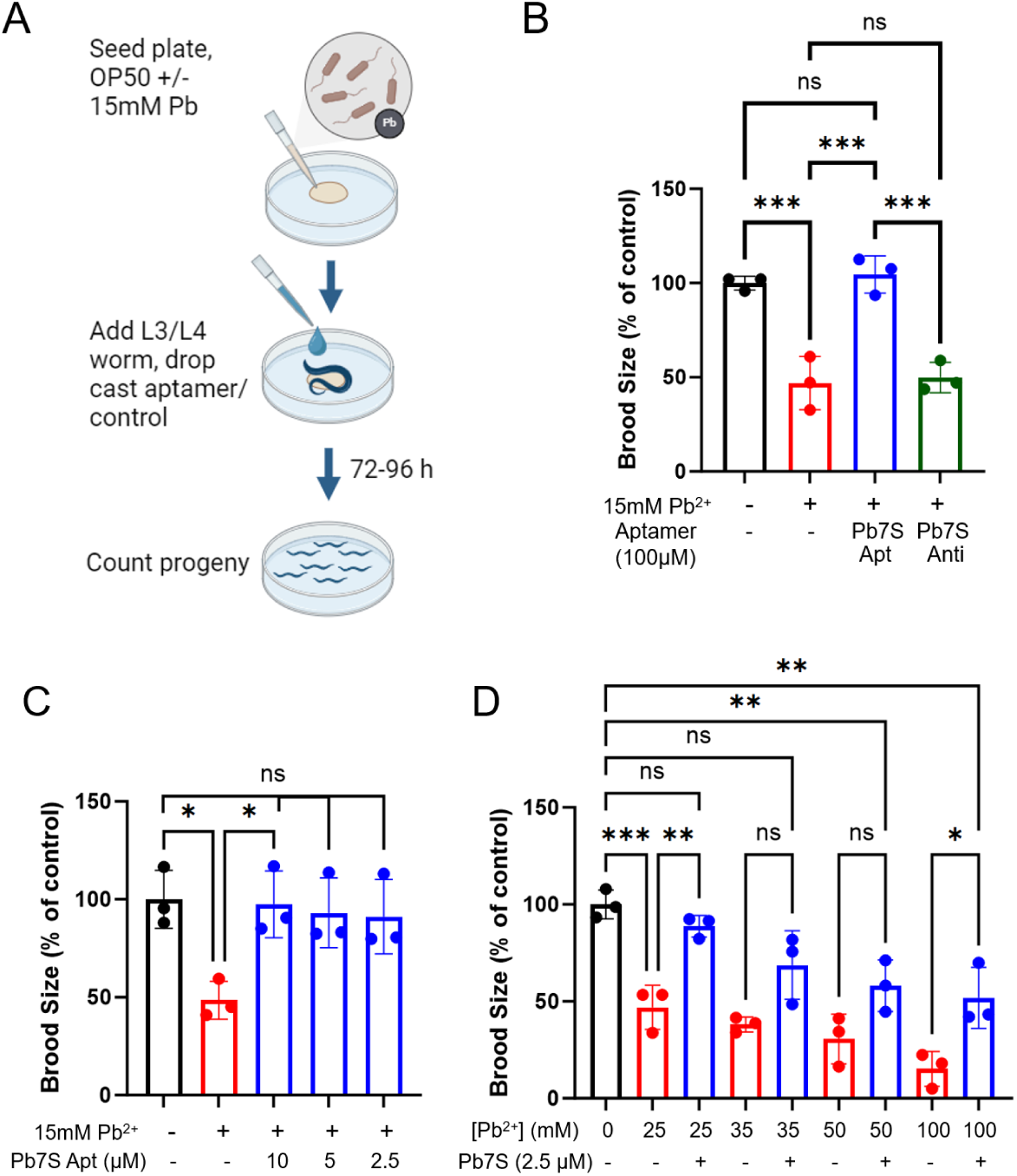
Pb7S ssDNA aptamers protect C. elegans from reproductive toxicity caused by lead. A) Schematic overview of the drop casting experimental method. B) Brood size assay. L3/L4 animals were fed 15 mM lead acetate mixed with OP50. ssDNA aptamers or water control were drop cast onto animals as indicated at 100 µM. Brood size was assessed by counting total offspring at 96 h. Apt, aptamer. Anti, antisense. One-way ANOVA P<0.0001. C) Titration of the Pb7S aptamer. ssDNA aptamers were applied using the drop cast method, and brood sizes determined at 96 h. One-way ANOVA P=0.0149. D) Titration of lead concentration. Lead acetate was mixed with OP50 at the indicated concentrations, then 2.5 µM of Pb7S was drop cast onto animals. Brood sizes were determined at 96 h. One-way ANOVA P<0.0001. Data in all panels are presented as the mean of three experimental replicates (n=3), wherein each overlaid point represents the experimental average of three technical replicates. Error bars are +/- S.E.M. *** P<0.001, ** P<0.005, * P<0.05, ns not significant, by one-way ANOVA with Tukey’s post-hoc tests for multiple comparisons.

By all three exposure methods, we find exposure to the Pb7S aptamer, but not the antisense (reverse complement) aptamer or scrambled controls, results in a normal brood size (Supplementary Fig. S5, S6, Fig. 2B). To further confirm these results, we examined the original longer form of the Pb7S aptamer (known as Pb7) as well as the long and short forms of a second lead-binding aptamer identified by known as Pb14^29^, using the soaking method. Both the Pb7 and Pb14 aptamers, in their long and short forms, were effective in preventing the lead-associated decrease in brood size using the soaking method (Supplementary Fig. S6). In these soaking experiments, a modest but significant protective effect of the antisense strands was observed. We observed no effect of the Pb7, Pb7S, Pb14 or Pb14S aptamers, or their antisense sequences, on the reproductive toxicity of manganese^41,42^ (Supplementary Fig. S6). The drop casing method proved to be more robust than feeding, and less toxic to the animals than soaking, so we continued with that method.

To thoroughly examine the protective effect of the Pb7S aptamer, we tested the aptamer at a range of both aptamer and lead concentrations using the drop casting method. The minimum effective concentration of aptamer required to achieve full protection from exposure at 15mM lead acetate was 2.5 µM (Fig. 2C). At 2.5 µM treatment, significant protection of animals was observed up to 100 mM lead acetate (Fig. 2D). Therefore, we have demonstrated the lead-specific, dose-dependent protection of animals from ingested lead toxicity by exposure to lead-binding ssDNA aptamers using multiple application methods.

Our feeding and soaking experiments determined there was no effect of the aptamers on manganese toxicity (Supplementary Fig. S5, S6). Manganese is a divalent cation and an essential trace element with well-established biological roles^43^; cadmium, like lead, is a divalent cation with no known biological role and has similarly been strongly correlated with cardiovascular disease risk^44^. To examine the specificity of the Pb7S aptamer in protecting *C. elegans* from reproductive lead toxicity, we determined that feeding of 10 mM cadmium caused an ∼50% decrease in brood size (Supplementary Fig. S7), then tested the ability of the lead-binding Pb7S aptamer to rescue the cadmium-induced reproductive toxicity. The Pb7S aptamer had no effect on cadmium-induced reproductive toxicity (Fig. 3A). Several cadmium-binding aptamers have been described in the literature^45,46^, with nanomolar affinities to cadmium (34.5 nM for Cd-4^46^). When the Cd-4 (Fig. 3B) or Cd2-2 (Supplementary Fig. S8) aptamers were applied in our brood size assay, we found no effect on the reproductive toxicity caused by lead. We conclude that the lead-binding aptamers are specific for lead, as they had no effect on the reproductive toxicity of manganese or cadmium, in addition to published evidence that the aptamers are highly specific for their target ions^29,45,46^.

**Figure 3.**
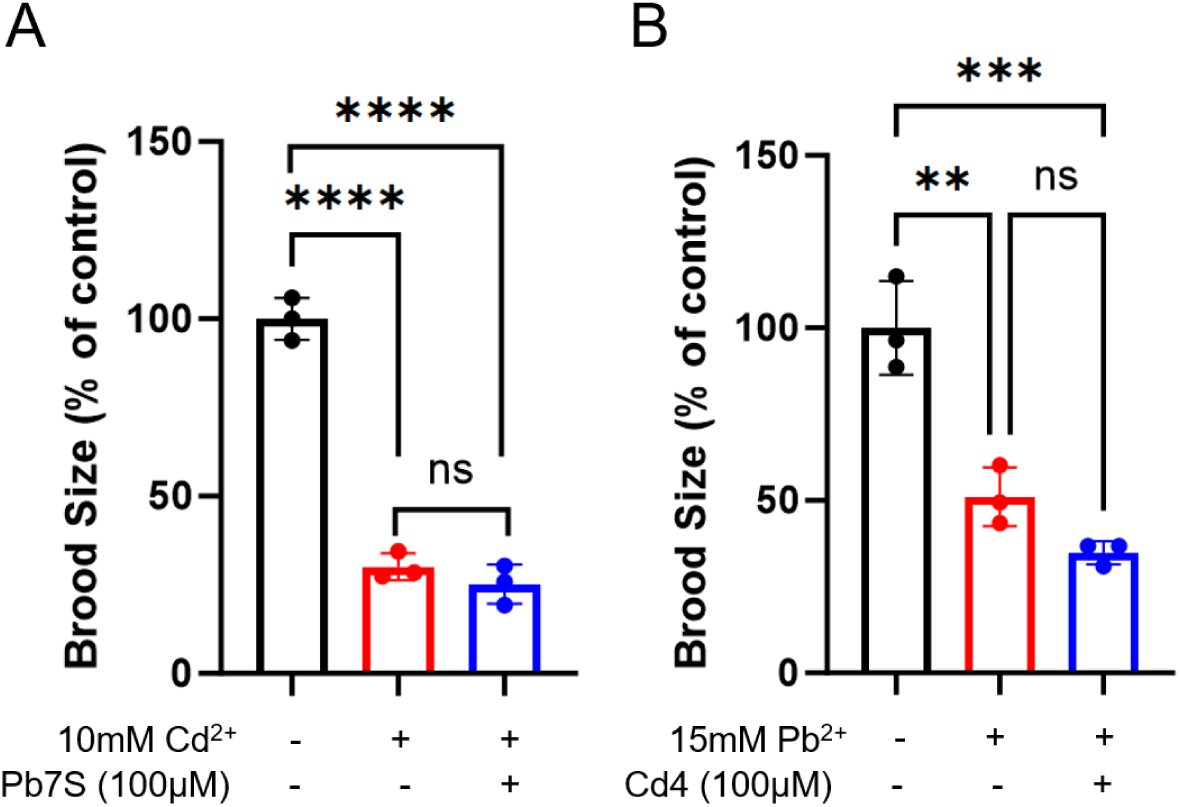
Pb7S aptamer is specific for lead and does not protect against cadmium toxicity. A) The Pb7S aptamer does not protect *C. elegans* from cadmium toxicity. 10mM cadmium chloride was mixed with OP50, and water or Pb7S aptamer was drop cast onto animals as indicated, then brood size assays were performed as described. One-way ANOVA P<0.0001. B) The cadmium-binding Cd4 aptamer does not protect *C. elegans* from lead toxicity. 15 mM lead acetate was mixed with OP50, and the Cd4 aptamer or water control was drop cast onto animals as indicated, then brood size assays were performed as described. One-way ANOVA P=0.0004. All data are presented as the mean of three experimental replicates (n=3), wherein each overlaid point represents the experimental average of three technical replicates. Error bars are +/- S.E.M. **** P<0.0001, *** P<0.001, ** P<0.005, ns not significant, by one factor ANOVA with Tukey’s post-hoc tests for multiple comparisons.

### Behavioral Toxicity of Lead is Prevented by Pb-Binding ssDNA Aptamers

Early lead exposure in children is well established to result in developmental neurotoxicity^3^. We therefore sought to employ a model of developmental neurotoxicity in the form of a behavioral assay in our *C. elegans* model. *C. elegans* are known to move away from aversive cues, a pattern of behavior known as avoidance^47^. To determine whether lead exposure negatively impacted *C. elegans* avoidance behavior during larval development, we exposed L1 stage worms to 15 mM lead acetate and allowed them to develop to the L3/L4 stage in the presence of lead, then tested their avoidance of three known noxious cues, all of which are known to require the function of the ASH neurons^48^; copper, glycerol, and quinine. In all cases, lead exposure during larval development resulted in a dampened avoidance response to all noxious cues (Fig. 4A-C, blue bars). Exposure to the aptamer without lead did not affect the normal avoidance behavior. When the animals were exposed to the Pb7S aptamer in addition to the lead, we observed a restoration of the normal avoidance behavior. Animals in all experiments exposed to a solvent control were not responsive, as expected (Fig. 4A-C, black bars). These results suggest that the aptamer protects the animals from lead-induced behavioral toxicity during development.

**Figure 4.**
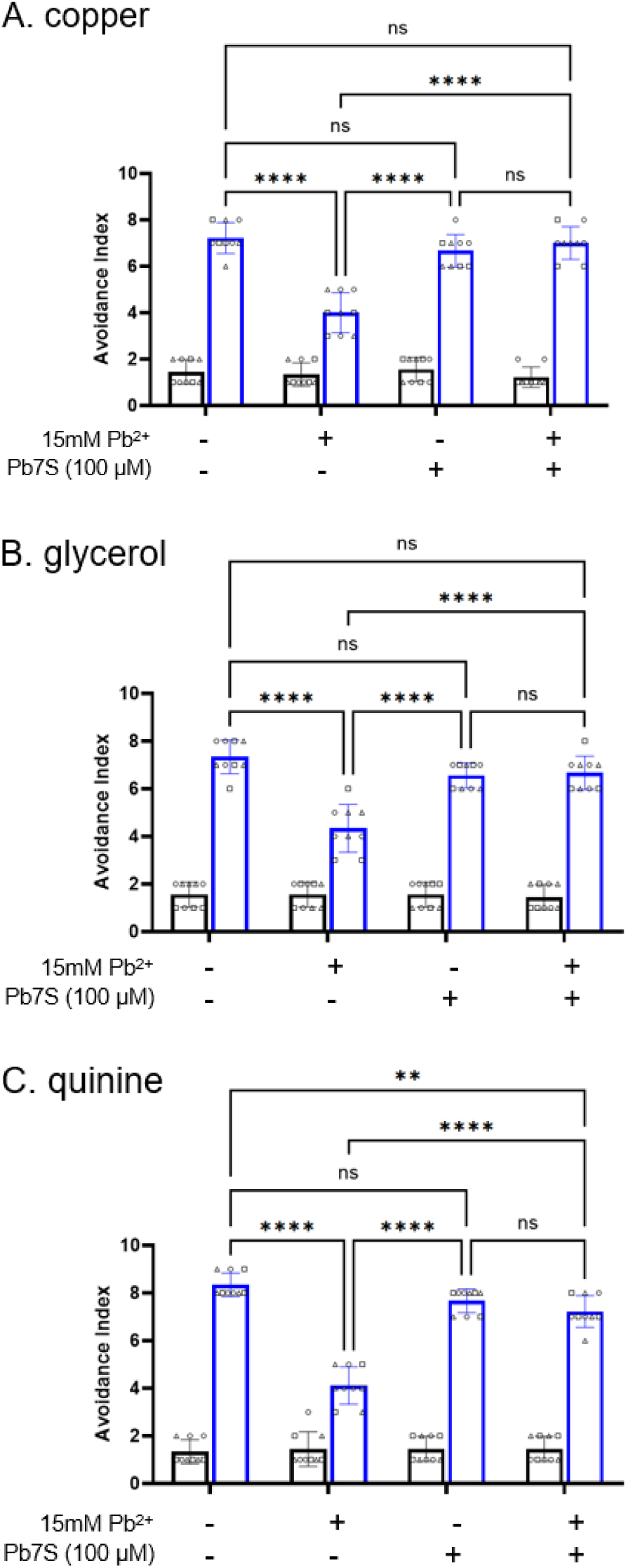
The Pb7S DNA aptamer protects C. elegans from developmental lead neurotoxicity. Animals were exposed to noxious chemical cues copper chloride (A), glycerol (B) and quinine (C) (blue bars) or a water solvent control (black bars) and observed for reversal of forward motion as described in Methods. For all panels, P<0.0001 by two-way ANOVA. Overlaid points indicate 9 biological replicates, containing 10 animals tested per replicate, for a total of 90 animals tested. Replicates tested on the same day are indicated with matching symbols. N=9. Error bars are +/- S.E.M. **** P<0.0001, ** P<0.005, ns not significant, by two-way ANOVA with Tukey’s post-hoc tests for multiple comparisons.

### Reproductive Toxicity of Lead is Prevented by Pb-Binding ssRNA Aptamers

Modified RNAs (siRNAs and mRNAs) have been approved in the U.S. for therapeutic and prophylactic uses^49^ and are a promising treatment modality^26,27^. To determine whether an RNA version of the Pb7S aptamer could also efficiently protect *C. elegans* from reproductive toxicity, we repeated our brood size assays using RNA versions of Pb7S and scrambled controls using the drop casting method. RNA aptamers result in protection of animals from lead-induced reproductive toxicity, whereas scrambled and antisense controls have no effect on brood size (Fig. 5A). To examine the protective range of the RNA Pb7S aptamer, we tested the aptamer at a range of both aptamer and lead concentrations. The minimum effective concentration of the RNA aptamer required to achieve full protection from exposure at 15 mM lead acetate was 2.5 µM (Fig. 5B). At 2.5 µM treatment, significant protection of animals was observed up to 100 mM lead acetate (Fig. 5C). These ranges were identical to those revealed in our ssDNA Pb7S aptamer testing (Fig. 2C, 2D). We conclude that ssRNA Pb7S aptamers are equally as effective as ssDNA aptamers in protecting *C. elegans* from reproductive toxicity.

**Figure 5.**
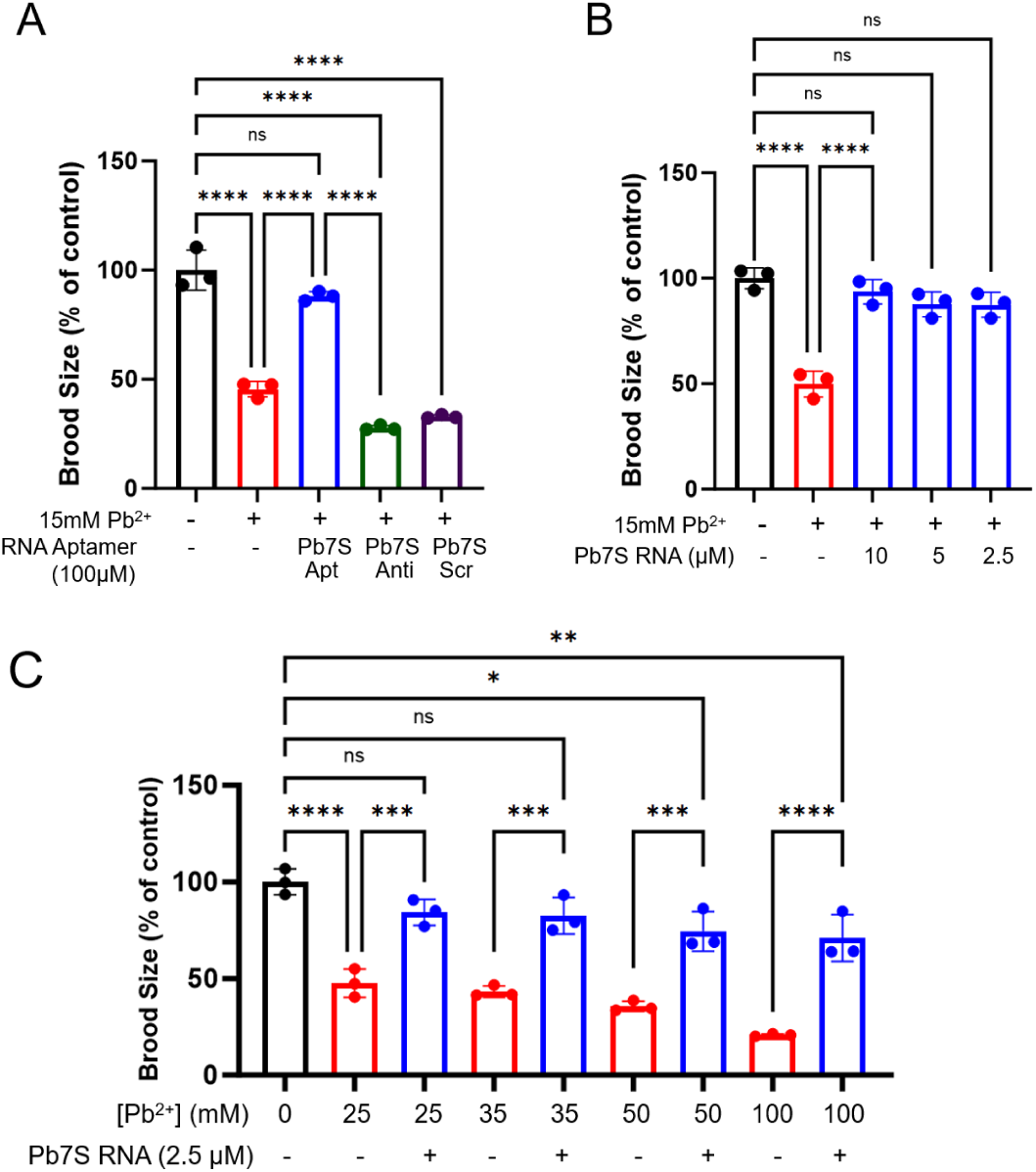
Pb7S ssRNA aptamers protect *C. elegans* from reproductive toxicity. A) Brood size assay. L3/L4 animals were fed 15mM lead acetate mixed with OP50. ssRNA aptamers or water control were drop cast onto animals as indicated at 100 uM. Brood size was assessed by counting total offspring at 96 h. Apt, aptamer. Anti, antisense. Scr, scrambled control. One-way ANOVA P<0.0001. B) Titration of the Pb7S ssRNA aptamer. Aptamers were applied using the drop cast method, and brood sizes determined at 96 h. One-way ANOVA P<0.0001. C) Titration of lead concentration. Lead acetate was mixed with OP50 at the indicated concentrations, then 2.5 uM of Pb7S RNA was drop cast onto animals. Brood sizes were determined at 96 h. One-way ANOVA P<0.0001. Data in all panels are presented as the mean of three experimental replicates (n=3), wherein each overlaid point represents the experimental average of three technical replicates. Error bars are +/- S.E.M. **** P<0.0001, *** P<0.001, ** P<0.005, * P<0.05, ns not significant, by one-way ANOVA with Tukey’s post-hoc tests for multiple comparisons.

### Toxicity of Lead in Cultured Cells is Prevented by Pb-Binding ssDNA Aptamers

To determine whether ssDNA Pb7S aptamer could protect mammalian cells from lead toxicity, we used cell proliferation assays to measure the effect of lead on cultured cell growth. We investigated the effects of lead exposure on cell proliferation in HEK293 cells and primary murine calvarial osteoblasts over a concentration range from 0 to 50 μM, and assessed the ability of aptamers to mitigate lead toxicity. Calvarial osteoblasts were isolated from calvaria of wildtype neonates at postnatal day 5. We observed a substantial and dose-dependent reduction in cell proliferation at 48 h of lead exposure in both cell types (Fig. 6A, 6B). Transfection of the Pb7S aptamer at concentrations of 10 nM or 50 nM partially or completely, respectively, protected the cells from cytotoxicity and preserved proliferation, relative to transfection of the scrambled control.

**Figure 6.**
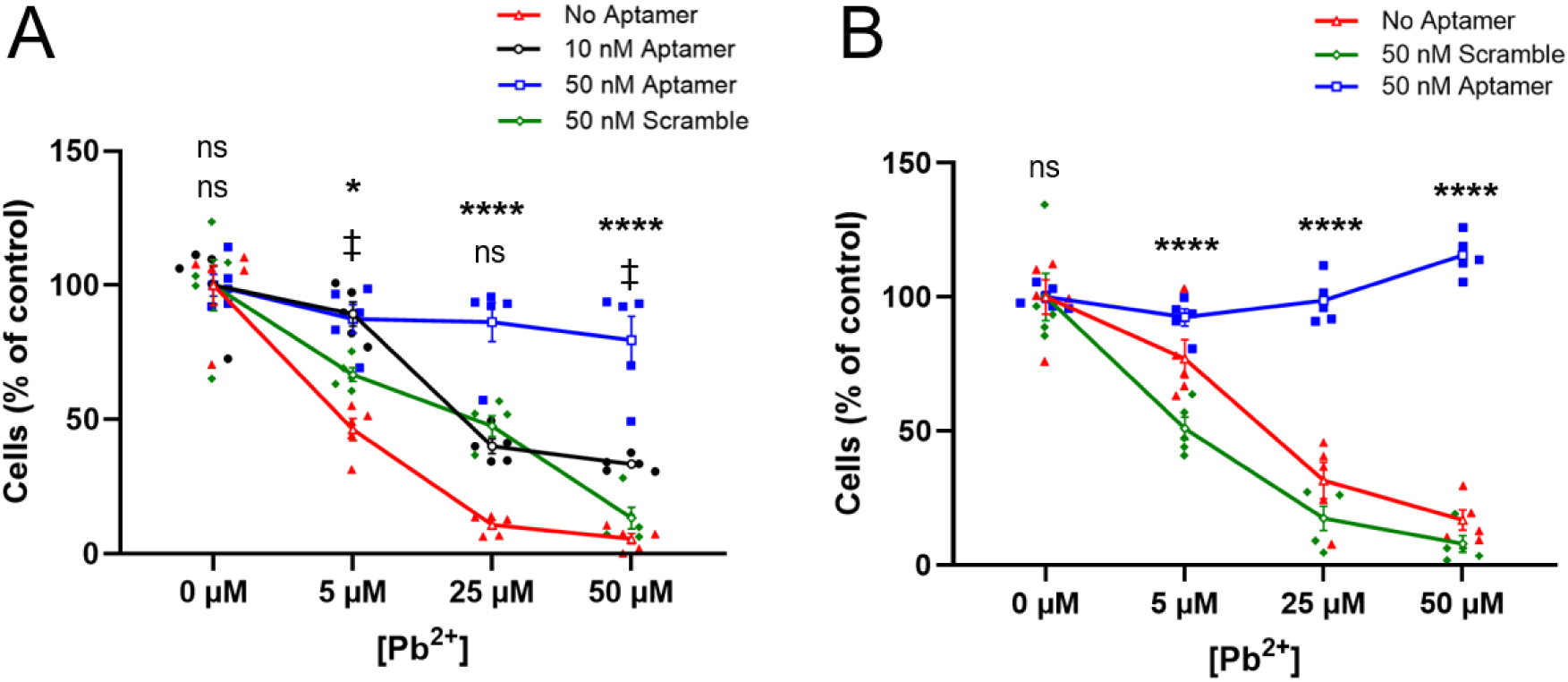
ssDNA Pb7S aptamers protect cultured HEK293T and primary calvarial osteoblasts from lead toxicity. HEK293 (A) or primary calvarial osteoblasts (B) were transfected with 10 nM (black line and symbols) or 50 nM (blue line and symbols) Pb7S aptamer, or 50 nM scrambled control (green line and symbols), or mock transfected (no aptamer, red line and symbols), then exposed to lead concentrations as indicated. P<0.0001 by two-way ANOVA for each panel. Data in both panels are presented as the mean of five experimental replicates (n=5), wherein each overlaid point represents the experimental average of 3-4 technical replicates. Bars are +/- S.E.M. **** P<0.0001, * P<0.05, for comparison between 50 nM Pb7S aptamer and 50 nM scramble conditions; ‡ P<0.05, for comparison between 10 nM Pb7S aptamer and 50 nM scramble control; ns not significant, by two-way ANOVA with Tukey’s post-hoc tests for multiple comparisons.

To establish the functional toxicity of lead in osteogenic differentiation of cultured primary calvarial osteoblasts, the expression of osteogenic marker genes, such as Runx2 (Fig. 7A), bone sialoprotein (Ibsp, Fig. 7B), type I collagen α 1 (Col1α1, Fig. 7C), and tissue-nonspecific alkaline phosphatase (Tnalp, Fig. 7D) were examined^50,51^. Cells were treated with different concentrations of lead over a 6-day period, with or without the transfection of the Pb7S aptamer, prior to RNA extraction and analysis by qPCR. We observed a significant inhibition of expression of these osteogenic genes (Fig. 7Ai – Di) in response to lead exposure. Notably, when cells were first transfected with 10 nM Pb7S aptamer (Fig. 7Aii-Dii), gene expression was not affected by the exposure to lead, effectively rescuing the expression of these genes from the inhibitory effects of lead.

**Figure 7.**
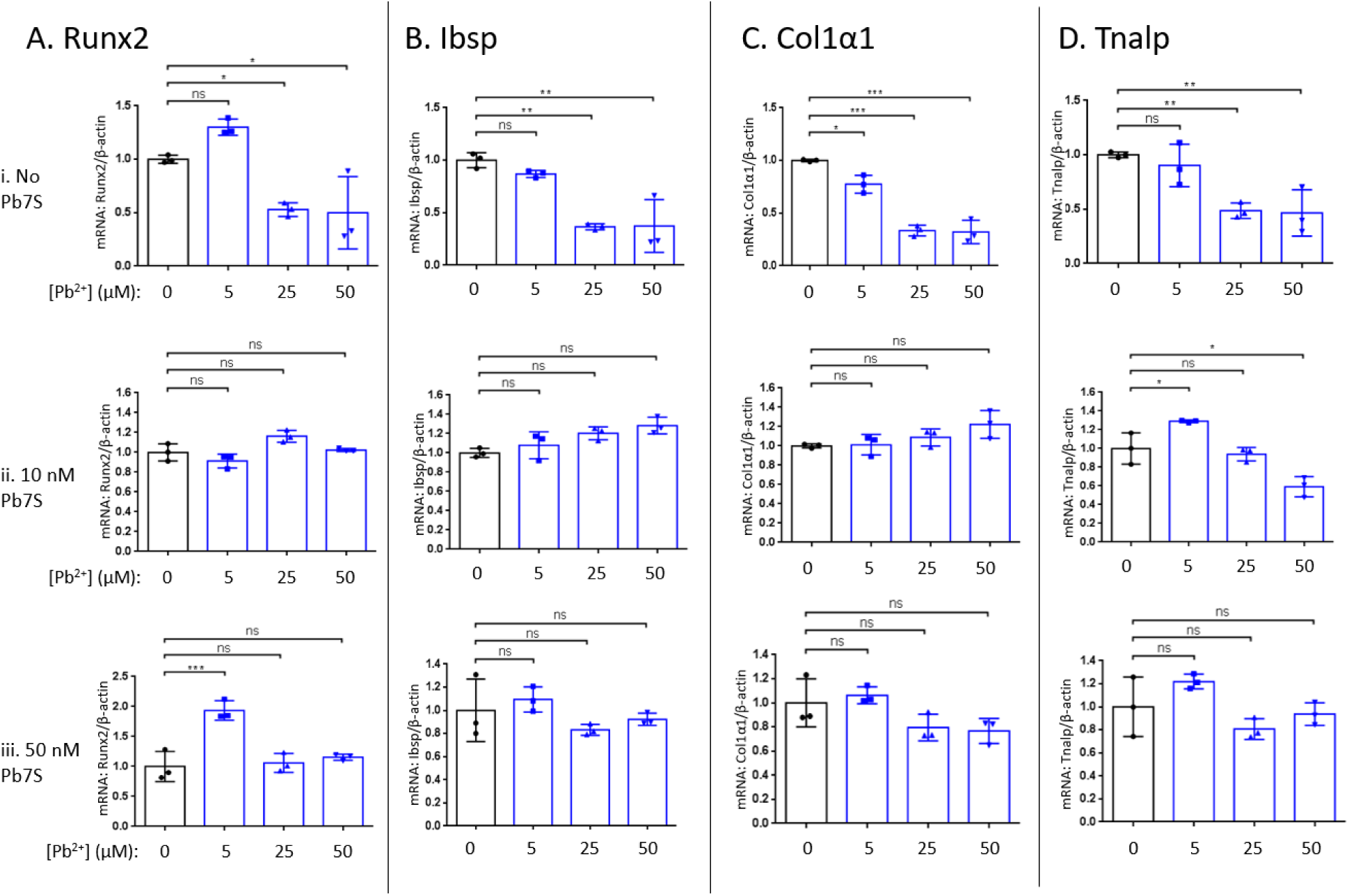
Changes in osteogenic gene expression caused by lead are prevented by the Pb7S aptamer. Primary calvarial osteoblasts were mock transfected (i) or transfected with the Pb7S DNA aptamer at 10 nM (ii) or 50 nM (iii), and treated with lead concentrations as indicated, then RNA was extracted and analyzed by qRT-PCR for osteogenic gene markers Runx2 (A), Ibsp (B), Col1a1 (C) or Tnalp (D). N=3. Error bars are +/- S.E.M. *** P<0.001, ** P<0.005, * P<0.05, ns not significant, by student’s t test versus 0 µM lead controls.

This protective effect was consistently replicated, and in the case of Runx2, enhanced, when cells were transfected with a higher aptamer concentration of 50 nM (Fig. 7Aiii-Diii). These findings highlight the aptamer’s capacity to prevent the suppression of osteogenic gene expression caused by lead, and to counteract the negative impact of lead on these critical factors in osteoblast development and differentiation.

We next examined alkaline phosphatase (ALP) activity in primary calvarial osteoblasts, which is a hallmark of differentiated, active osteoblasts^52^. In alignment with the gene expression results, increasing lead exposure resulted in decreased ALP activity (Fig. 8), suggesting that lead exposure negatively impacts osteogenic differentiation. However, transfection of the ssDNA Pb7S aptamer effectively preserved ALP function and increased ALP staining. The results suggest that both osteoblast survival and function are protected from the toxic effects of lead by the presence of the aptamer.

**Figure 8.**
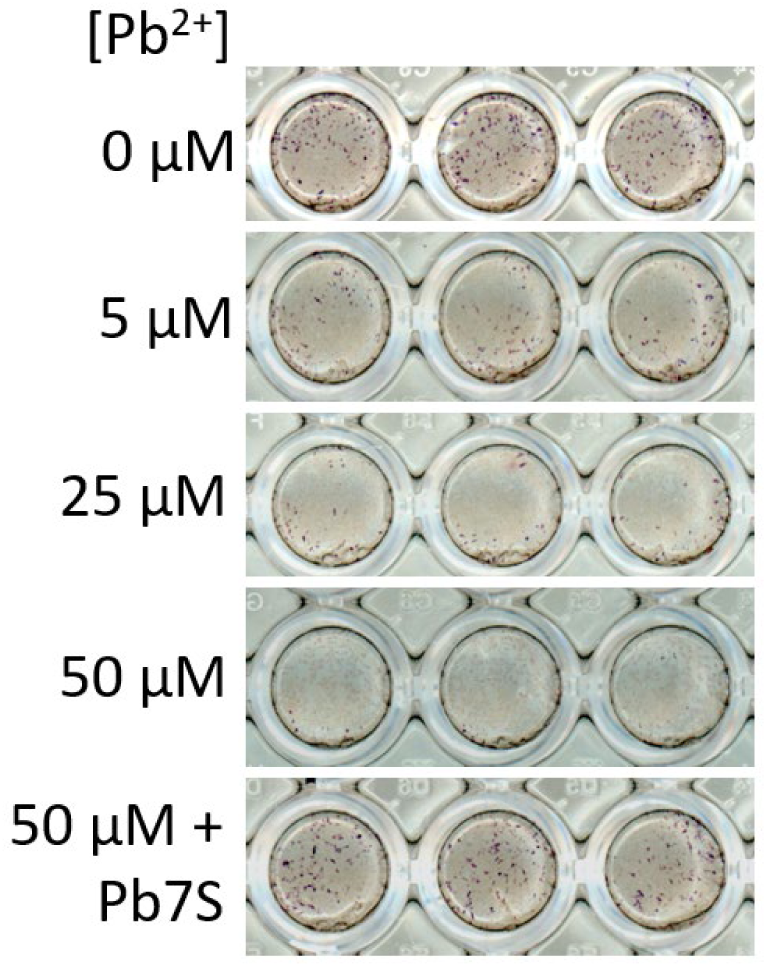
Lead induced decrease in osteogenic differentiation is prevented by exposure to the ssDNA Pb7S aptamer. Primary murine calvarial osteoblasts were transfected with Pb7S DNA aptamer as described, or a mock transfection control, then exposed to lead acetate concentrations as indicated. ALP activity was assessed by Fast Blue staining. Wells shown are technical replicates.

## DISCUSSION

The current standard of treatment for lead toxicity is chelation therapy with oral medication, or EDTA chelation by intravenous administration^53^. However, these therapies are typically only offered in the case of extremely high lead levels (blood lead levels above 45 µg/dL for children and 70 µg/dL for adults^53^), despite the fact that much lower levels are associated with negative health consequences, as discussed above^18,19^. The recommended course of action for lower blood lead levels (3.5 – 45 µg/dL) is to continue to monitor the lead levels of the patient and attempt to identify and eliminate the source of contamination^53^. Again, this course of action cannot reverse permanent neurologic damage, nor can it prevent the accumulation of lead in bones.

Many lead exposures are via ingestion from environmental sources: water, food, or incidental ingestion of contaminated dust or paint chips^53^. When the water or working conditions are to blame, it may be difficult or impossible to eliminate lead from the environment completely. This is particularly true for low-income families. Analysis of the NHANES data from 2015-2018 revealed that Hispanic, Black, and low-income children of all races (ages 1-5) were significantly more likely to have elevated blood lead levels as compared to the national average of all races/incomes^20^. The neurotoxic effects of lead are permanent, leading to life-long cognitive deficits in these children and creating a disease burden that is borne disproportionately by racially diverse and low-income communities^54^. Therefore, there is not only an urgent need but an environmental justice obligation to develop accessible and cost-effective methods to protect people from lead.

In this work, we demonstrate that the Pb7S aptamer binds with sensitivity and specificity to lead ions, and can efficiently chelate these ions to inhibit their toxicity. ssDNA aptamers applied to *C. elegans* protected animals from reproductive toxicity and behavioral toxicity caused by lead ingestion. The aptamers are specific to lead, and do not affect cadmium toxicity or manganese toxicity. The protective effect of Pb7S was observed with three different methods of aptamer delivery: feeding, soaking, and drop casting. Our CD spectroscopy experiments, in alignment with the literature^30,38^, suggest that Pb7S forms a stable complex with lead ions, and that there is no discernible interference of the interaction by calcium. The dramatic shift in the CD spectra of lead in complex with Pb7S supports the possibility that the aptamer chelates lead in a stable G4 structure, though additional investigation will be needed to determine the specific structure and binding kinetics of the complex. Finally, exposure of *C. elegans* to the Pb7S aptamer is protective against lead toxicity. However, future studies are needed to examine the protection mechanism, including ascertaining intracellular lead levels and translation of the effects to whole vertebrate animals.

Our studies revealed that RNA versions of the Pb7S aptamer were equally as effective as DNA aptamers in providing a protective effect in our reproductive toxicity assays. This raises the possibility that RNA-based aptamers could be developed for clinical use as an alternative to intravenous sodium edetate (EDTA) therapy for lead poisoning. These aptamer-based chelation over chemical chelators like EDTA could improve specificity for lead ions leading to decreased calcium excretion^55^, as well as administration (a subcutaneous injection^56,57^ at home versus intravenous therapy in a clinic). Modified RNAs are increasingly being used as prophylactics and therapeutics in the form of mRNA vaccines, antisense oligonucleotides (ASOs), and small interfering (si)RNAs, and their impact is expected to increase rapidly and significantly^49^. Aptamers are known from human studies to be generally safe and efficiently excreted by the kidneys^58^, which is a drawback for ASOs or siRNAs used therapeutically but in the context of eliminating lead ions or other toxins may be an asset.

In human HEK293 cells and murine calvarial osteoblasts, we demonstrated the protective effects of the Pb7S aptamer on cell proliferation, osteogenic gene expression, and osteogenic differentiation. In a rat model of developmental lead exposure, dimercaptosuccinic acid (DMSA) chelation partially reversed negative changes in bone microarchitecture and osteogenic markers associated with lead exposures^59^, which suggests that the decreased osteogenic potential associated with lead toxicity may be reversible with chelation. Our results suggest therapeutic potential of aptamers to protect cells and organ systems from lead toxicity. Whether aptamers could be used to reverse some of its toxic effects is unknown, but is an important future avenue of investigation.

A 10-year, large scale clinical trial (the Trial to Assess Chelation Therapy, TACT) studied the use of intravenous chelation therapy with EDTA to reduce the incidence of repeat adverse events and death in patients that had already suffered one heart myocardial infarction (MI)^60^. The study found that heavy metal chelation resulted in an 18% decrease in repeat adverse CVD events. In diabetic patients, EDTA chelation reduced repeat CVD events by 41%^61^. Importantly, the study participants were not selected based on blood lead levels; they were patients who had MI previously. The implication is that our standard lifetime exposures to metals, lead and cadmium in particular^4^, are increasing our risk of heart disease, and that decreasing these metals in our system results in a decreased risk of CVD and death. Aptamer-based chelation strategies will require further research before they are ready for the clinic. However, given that over 14% of the U.S. population is estimated to be diabetic^62^ and that CVD is the number one leading cause of death in the U.S.^63^, the potential impact of decreasing toxic heavy metal absorption, at least in part by employing aptamers, cannot be overstated.

## MATERIALS AND METHODS

### Reagents, cell lines, and materials

Aptamer sequences are listed in Supplementary Table S1. All aptamers were synthesized as standard desalted single-stranded DNA or RNA oligonucleotides by IDT-DNA (Coralville, IA, and Research Triangle Park, NC, USA). Wildtype N2 *C. elegans* were obtained from the Caenorhabditis Genetics Center (University of Minnesota, MN), and were used for all experiments. N2 animals were maintained on nematode growth media (NGM) plates and fed with OP50 *E. coli*. HEK293 cells were obtained from the American Type Culture Collection in Rockville, Maryland. These cells were cultured in Dulbecco’s Modified Eagle Medium (DMEM) from Corning, with the addition of 10% Fetal Bovine Serum (FBS), 2 mM L-glutamine, 1% penicillin/streptomycin, and 1% nonessential amino acids, all of which were also provided by Corning. The cells were maintained at 37°C in a humidified environment with 5% CO2.

### Primary murine calvarial osteoblast cell culture

Primary calvarial osteoblasts were isolated from calvaria of 5-day-old neonates using collagenase type II (50 mg/ml, Worthington, LS004176)/dispase II (100 mg/ml, Roche, 10165859001). Osteoblasts were maintained in alpha-minimal essential medium (aMEM) (Gibco) containing 10% FBS (Gibco), 2mM L-glutamine (Corning), 1% penicillin/streptomycin (Corning), and 1% nonessential amino acids (Corning) and differentiated with ascorbic acid (200 uM, Sigma, A8960). and β-glycerophosphate (10 mM, Sigma, G9422). To induce osteogenic differentiation, the growth media was supplemented with ascorbic acid (200 mM, Sigma, A8960) and beta-glycerophosphate (10 mM, Sigma, G9422).

### Osteoblast differentiation analysis

To assess osteoblast differentiation, alkaline phosphatase (ALP) activity was determined as described previously^52^. In brief, differentiated osteoblasts were washed with phosphate-buffered saline (PBS) and incubated with a solution containing 6.5 mM Na_2_CO_3_, 18.5 mM NaHCO3, 2 mM MgCl_2_, and a phosphatase substrate (Sigma, S0942). ALP activity was measured using a spectrophotometer. ALP staining was performed using Fast Blue (Sigma, FBS25) and Naphthol AS-MX (Sigma, 855) after fixation in 10% neutral formalin buffer.^63^

### Lead treatment and aptamer transfection

Aptamer transfections were conducted using Lipofectamine RNAi MAX reagent in Opti-MEM®I Reduced Serum Medium (from Invitrogen, Carlsbad, CA, USA) following the manufacturer’s protocol. Calvarial osteoblasts were plated at a density of 5x10^4 cells per well in a 12-well plate. One day later, the cells were exposed to either lead acetate alone at indicated concentrations, or a combination of lead and aptamers (10nM, 25nM and 50nM) in Opti-MEM (reduced serum medium) using Lipofectamine RNAi MAX reagent from Invitrogen for 6 hours. After 6 hours, the culture medium was changed to a differentiation medium for a period of 6 days. The differentiation medium was changed every 2 days. Cells were harvested for RNA isolation after 6 days of differentiation.

### Cell proliferation assays

The Alamar Blue assay was performed in 96-well plates to assess cell proliferation according to the manufacturer’s instructions. Each well was initially seeded with a cell density ranging from 5,000 to 10,000 cells. After a 24-hour incubation, the cells were exposed to various concentrations of Pb (0, 0.25uM, 0.5uM, 1uM, 5uM, 10uM, 25uM and 50uM). Concurrently, the cells were transfected with three different concentrations of Aptamers (Apt) at 10nM,25nM and 50nM. These treatments were applied for both 24 and 48 hours. The culture medium was aspirated, and the cells were rinsed with phosphate-buffered saline (PBS). Then, cells were incubated with 100ul of tenfold diluted Alamar Blue solution (Invitrogen, DAL1100). After a 3-hour incubation period, the fluorescence of the Alamar Blue was measured at excitation and emission wavelengths of 575 and 595 nm using a Tecan Genios microplate reader. Measurements of cell viability using the Alamar Blue assay were taken 24 hours after the initial setup.

### Quantitative RT-PCR analysis

Total RNA was isolated from cells using QIAzol (QIAGEN) and cDNA was synthesized with 1ug of RNA using the High-Capacity cDNA Reverse Transcription Kit (Applied Biosystems). SYBR green chemistry was used for quantitative determination of the various mRNAs for Runx-2, Ibsp, Col1α1, Tnalp, and a housekeeping gene β-actin. Quantitative RT-PCR was performed using SYBR® Green PCR Master Mix (Bio-Rad) with CFX connect RT-PCR detection system (Bio-Rad).

### Brood Size Assays

NGM plates were seeded with 75 µL of *E. coli* OP50, mixed with or without lead acetate concentrations as indicated (and/or ssDNA aptamers as indicated for the feeding method). One L3-L4 N2 wildtype aged-synchronized worm was plated onto each seeded plate. For the drop casting method, 10 µL of aptamer solution (ssDNA or RNA) at the indicated concentration was applied to the animal by pipetting after plating. The brood size was determined at 72 or 96 hours as indicated.

### Avoidance Assay

Assays were performed as described previously^48,64^. Briefly, developmental plates containing 20 L1 N2 aged-synchronized worms were plated to NGM plates with or without 15 mM lead acetate, and with or without drop casting of 100 µM Pb7S ssDNA aptamer, and incubated at 20°C for 72 hours where they developed to the L3/L4 stage. *C. elegans* from the developmental plate were transferred to an acclimation plate in the testing room and allowed to acclimate to the new plate for 30 minutes. A drop of either water (solvent control) or stimulus (1 M glycerol,10 mM Copper (II) chloride dihydrate, 1 mM quinine) was placed on the tail of forward moving animals, and their response was scored as either an avoidance response (two body bends of reverse motion), or no response. The total number of avoidances was divided by the total number of drops to generate an avoidance index for that plate. This was repeated for at least 3 plates of 10 animals each, over at least three different days.

## STATISTICAL ANALYSIS

Statistical analyses were performed with GraphPad Prism software. All *C. elegans* and cell culture data were first analyzed by one-way or two-way ANOVA, as appropriate, followed by post-hoc testing as indicated in the figure legend. qRT-PCR results were analyzed by student’s t tests.

## SUPPLEMENTARY INFORMATION

Supplementary Figures 1 – 8, and Supplementary Table 1, are associated with this manuscript.

## DATA AVAILABILITY

The data that support the findings of this study are available from the corresponding author upon reasonable request.

## ACKNOWLEDGEMENTS

This work was supported in part by the generous philanthropy of Mr. Robert Ferrari of Northeast Water Solutions Inc. and Ferrari Engineering Inc. A.A. and S. Ramis de Ayreflor Reyes were supported in part by the Robert F. Ferrari ’74 Graduate Research Fund. N.G.F is supported in part by seed grant funding and new faculty start-up funds from WPI. We thank Caroline Muirhead and Elizabeth DiLoreto for technical assistance and advice. J.H.S. is supported by NIH/NIAMS (R01AR068983, R21AR073331), the International FOP Association, AAVAA Therapeutics, and Dong-A ST.

## AUTHOR CONTRIBUTIONS

A.A. and S. Ramis de Ayreflor Reyes performed all experiments with the following exceptions: A.A.J. performed HEK293 and murine osteoblast experiments; A.M.O. performed the cadmium toxicity curve; S. Reis performed behavioral toxicity on L3/L4 *C. elegans*; E.B. and N.G.F. performed CD spectroscopy experiments. Mentoring and supervision: J.H.S., A.A.S., J.S., N.G.F. Financial support: J.S. and N.G.F. Writing, original draft: N.G.F., A.A. and A.A.J. All authors contributed to editing and revision and have reviewed and approved the final manuscript.

## MATERIALS & CORRESPONDENCE

Correspondence and material requests should be addressed to N.G.F

