## Supplemental Figures, Tables, and Methods for "Nucleic Acid Aptamers Protect Against Lead (Pb(II)) Toxicity"

**Supplementary Table S1: Sequences of all aptamers used in this study.**

| Aptamer | Sequence |
|---|---|
| Pb7 aptamer | GGAGGCTCTCGGGACGACGGCA<br>GGGCTGTCGTACGGTTTGTCGAA<br>GGTGTCGTCCCGATGCTGCAATC<br>GTAAGAAT |
| Pb7 antisense | ATTCTTACGATTGCAGCATCGGGA<br>CGACACCTTCGACAAACCGTACG<br>ACAGCCCTGCCGTCGTCCCGAGA<br>GCCTCC |
| Pb7S aptamer | GGGACGACGGCAGGGCTGTCGTA<br>CGGTTTGTCGAAGGTGTCGTCCC<br>GA |
| Pb7S antisense | TCGGGACGACACCTTCGACAAAC<br>CGTACGACAGCCCTGCCGTCGTC<br>CC |
| Pb7S scramble | GCGGGCGATCTGCGGACGTTCTG<br>AGCCTGACTGAGTGGGGACGCTG<br>TA |
| Pb14 aptamer | GGAGGCTCTCGGGACGACGGCC<br>AGTAGCTGACATCAGTGTACGATC<br>TAGTCGTCCCGATGCTGCAATCGT<br>AAGAAT |
| Pb14 antisense | ATTCTTACGATTGCAGCATCGGGA<br>CGACTAGATCGTACACTGATGTCA<br>GCTACTGGCCGTCGTCCCGAGAG<br>CCTCC |
| Pb14S aptamer | TGACGACGGCCAGTAGCTGACAT<br>CAGTGTACGATCTAGTCGTCA |
| Pb14S antisense | TGACGACTAGATCGTACACTGATG<br>TCAGCTACTGGCCGTCGTCA |
| Non-specific scramble | AGAGGTTGGTTGCTGTGCGGACG<br>TACGGCCTGTCCGAAGCGCCGTG |
| 5’ fluor (FAM or YakYel) Pb7S | CTCAGTCGGGACGACGGCAGGG<br>CTGTCGTACGGTTTGTCGAAGGT<br>GTCGTC (underlined sequence base<br>pairs with quench strand) |
| Pb7S quench (with 3’ IAB, BHQ or DAB) | CGTCCCGACTGAG |

**Supplementary Table 2. Mouse primer sequences for qRT-PCR.**

| Name | Forward Primer | Reverse Primer |
|---|---|---|
| Runx2 | TACAAACCATACCCAGTCCCTGTTT | AGTGCTCTAACCACAGTCCATGCA |
| Ibsp (Bsp) | CAGGGAGGCAGTGACTCTTC | AGTGTGGAAAGTGTGGCGTT |
| Col1α1 | ACTGTCCCAACCCCCAAAG | ACGTATTCTTCCGGGCAGAA |
| Tnalp (Alp) | CACAATATCAAGGATATCGACGTGA | ACATCAGTTCTGTTCTTCGGGTACA |
| β-actin | GGCTGTATTCCCCTCCATCG | CCAGTTGGTAACAATGCCATGT |

### Supplementary Methods

*In vitro* fluorescence detection assays**.** To compare the different combinations of fluorescent molecule and quencher pairs, fluorescence detection assays were performed essentially as described [ref. 29]. Mixtures were prepared of fluorescent aptamers at 1X (0.1 μM) and quench strands at 6X (0.6 μM), in SELEX buffer (20 mM Tris-HCl, 10 mM sodium acetate, pH = 7.4) with lead nitrate in concentrations of 0, 1, 2.5, 5 ,10, and 20 μM. The plates incubated at room temperature for an hour, and a Perkin Elmer Victor3 plate reader was used to read green fluorescence (ex. 485, em. 515). Five experimental replicates were performed.

**Supplementary Figure S1. The aptamer-based fluorescence assay for Pb(II).** 5'FAM labeled aptamers are hybridized with short 3' DAB labeled quench strands to form a partial double helix at the 5' end of the aptamer. The proximity of the DAB to FAM quenches fluorescent signal. Upon addition of Pb(II), the FAM-aptamer dissociates from the quench strand and forms a complex with the lead ion, releasing the DAB-quench strand and resulting in fluorescence. The system is described in Chen et al., 2018 [ref. 29]. In some experiments as indicated, the fluor Yakima Yellow replaced FAM, and the quenchers Iowa Black (IAB) or Black Hole Quencher 1 (BHQ1) replaced DAB. Figure created with BioRender.com.

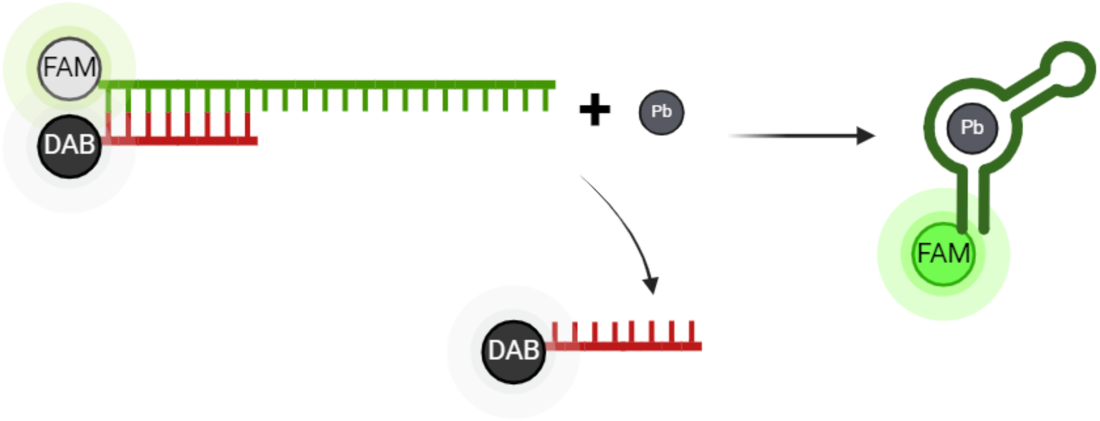

**Supplementary Figure S2. Performance of a fluorescence-based detection system for lead ions.** Fluors FAM (A) and Yakima Yellow (YakYel, B) were tested in combination with three quenchers, DAB, BHQ1 and IABk. A two-way ANOVA test and post hoc analysis determined there was no statistical difference from zero lead at any of the lead concentrations in the FAM-Pb7S data trials. In the YakYel-Pb7S trial (B), only the YakYel-Pb7S-DAB aptamer complex and YakYel-Pb7S-IABk complex had a statistical significance between the no lead and 20 μM lead concentrations with P values of 0.042 and 0.0415 respectively. Error bars are +/- S.E.M., n=5

A.

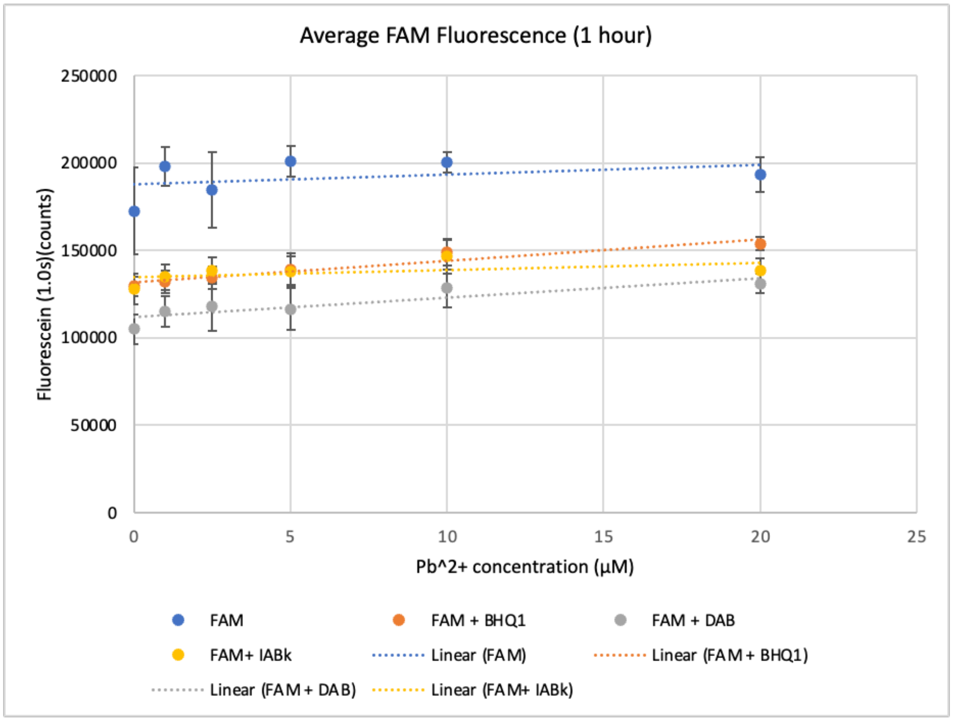


B.

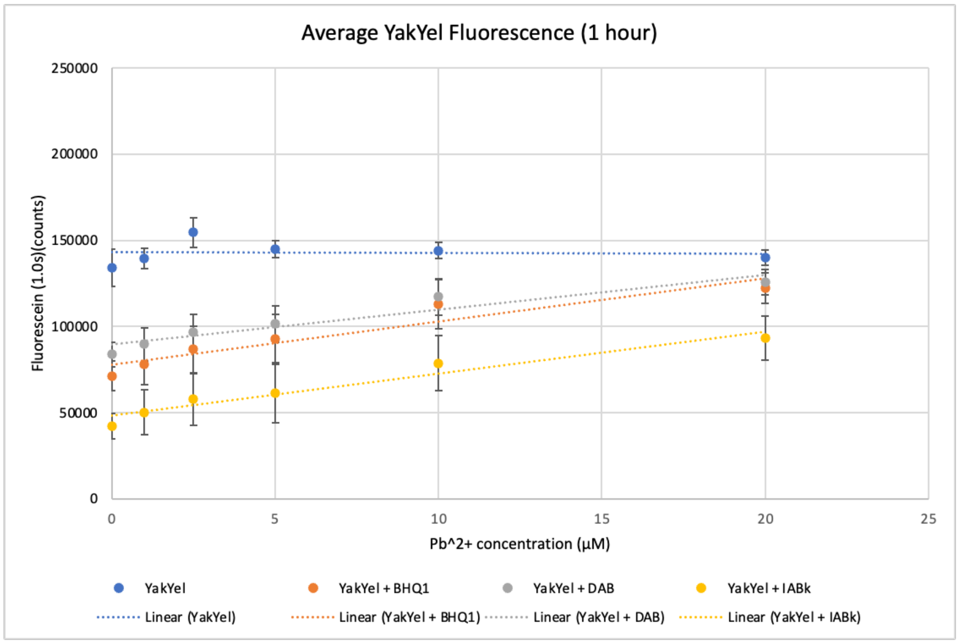

**Supplementary Figure S3. The effect of pH on the detection of lead by the fluorescence based Pb7S system**. A) The detection of lead by the YakYel-Pb7S aptamer and IAB quencher was determined at pHs ranging from 5.5 to 8.4. Error bars are +/- S.E.M., n=5. B) The slope of each line from part A, which is indicative of the dynamic range of detection, was plotted as a function of pH.

A.

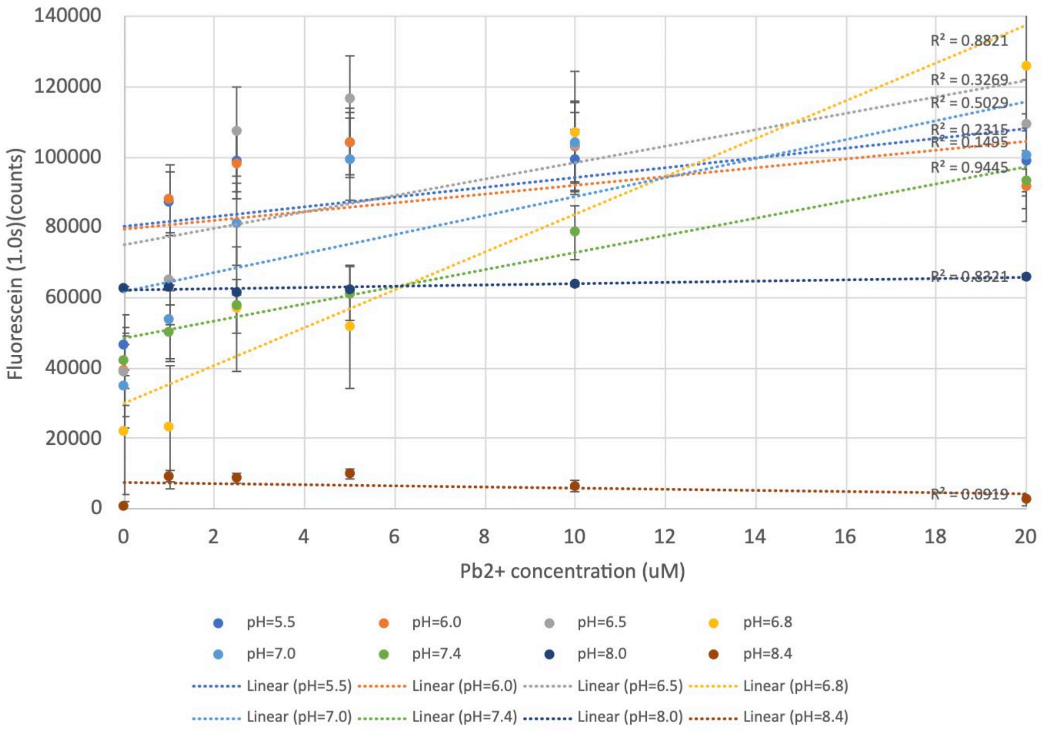


B.

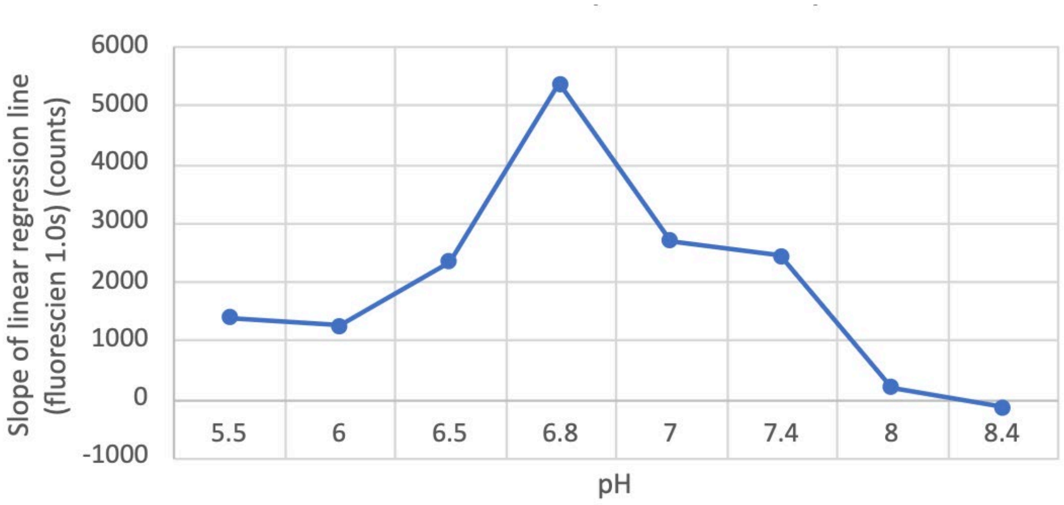

**Supplementary Figure S4. Determining the effect of lead concentration on brood size**. Lead concentrations indicated were mixed into 75μL of OP50 feed stock, plated to standard NGM plates, and allowed to dry. One L3/L4 animal was plated, and progeny were counted after 72 hours. Error bars are +/- S.E.M., n=3 replicates of the experiment. Each replicate at each concentration is the average of progeny from three animals.

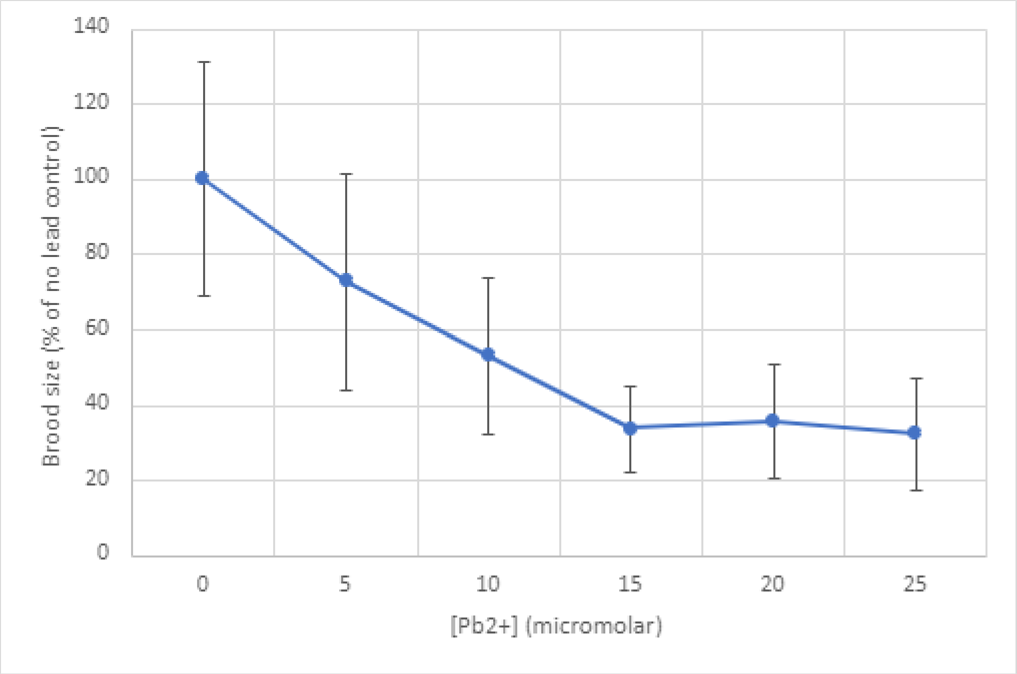

**Supplementary Figure S5. Feeding of lead-binding aptamers to *C. elegans* protects the animals from reproductive toxicity**. A) Schematic representation of the experimental workflow. Aptamers were mixed at 100μM into the OP50, with or without lead at 15mM. Then animals were plated to this mixture and offspring were counted after 96 hours. B) Brood size counts for *C. elegans* exposed to lead and aptamers by the feeding strategy. Feeding of the Pb7S aptamer, but not the antisense Pb7S, significantly increased brood size relative to the lead exposed control, whereas the aptamer has no effect on manganese toxicity. Error bars are +/- S.E.M., n=3. *=p<0.05.

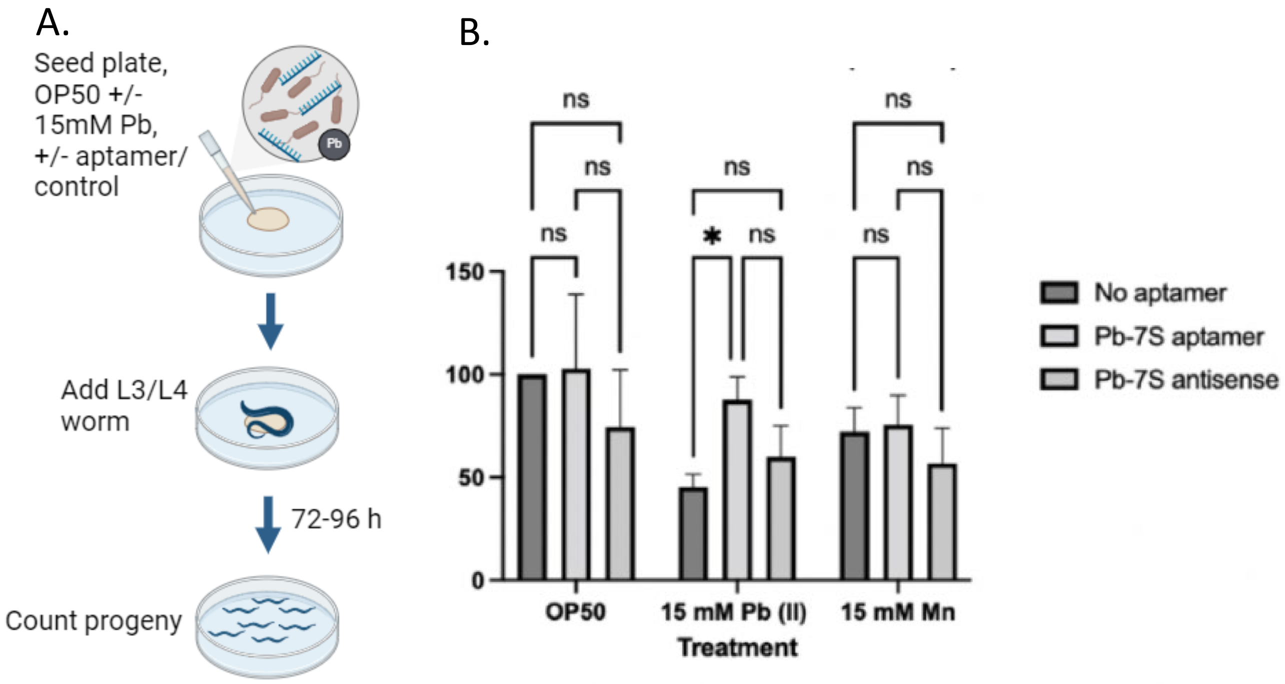

**Supplementary Figure S6. Soaking of *C. elegans* in lead-binding aptamers protects the animals from reproductive toxicity**. A) Schematic representation of the experimental workflow. Animals were soaked in M9 medium containing aptamers at 100µM for 2.5 h, then plated to NGM plates prepared with OP50, with or without lead at 15mM. Offspring were counted after 96 hours. B-E) Brood size counts for *C. elegans* exposed to lead and aptamers by the soaking strategy. Soaking in the Pb7 (B, ANOVA xxx), Pb7S (C, ANOVA xxxx), Pb14 (D, ANOVA xxx), or Pb14S (E, ANOVA xxx) aptamers, significantly increased brood size relative to the control exposed to lead and soaked in a Pb7S scrambled sequence (scramble). The antisense strands in some instances provided partial protection from lead toxicity. The aptamer and antisense strands have no effect on manganese toxicity. Error bars are +/- S.E.M., n=5. Water and scramble-soaked controls are identical for panels B-E; the aptamer treatments were separated into panels for ease of interpretation and statistical analysis. Results were analyzed by two-way ANOVA, followed by Tukey's post hoc tests for multiple comparisons. **** P<0.0001, *** P<0.001, ** P<0.005, *P<0.05, ns not significant.

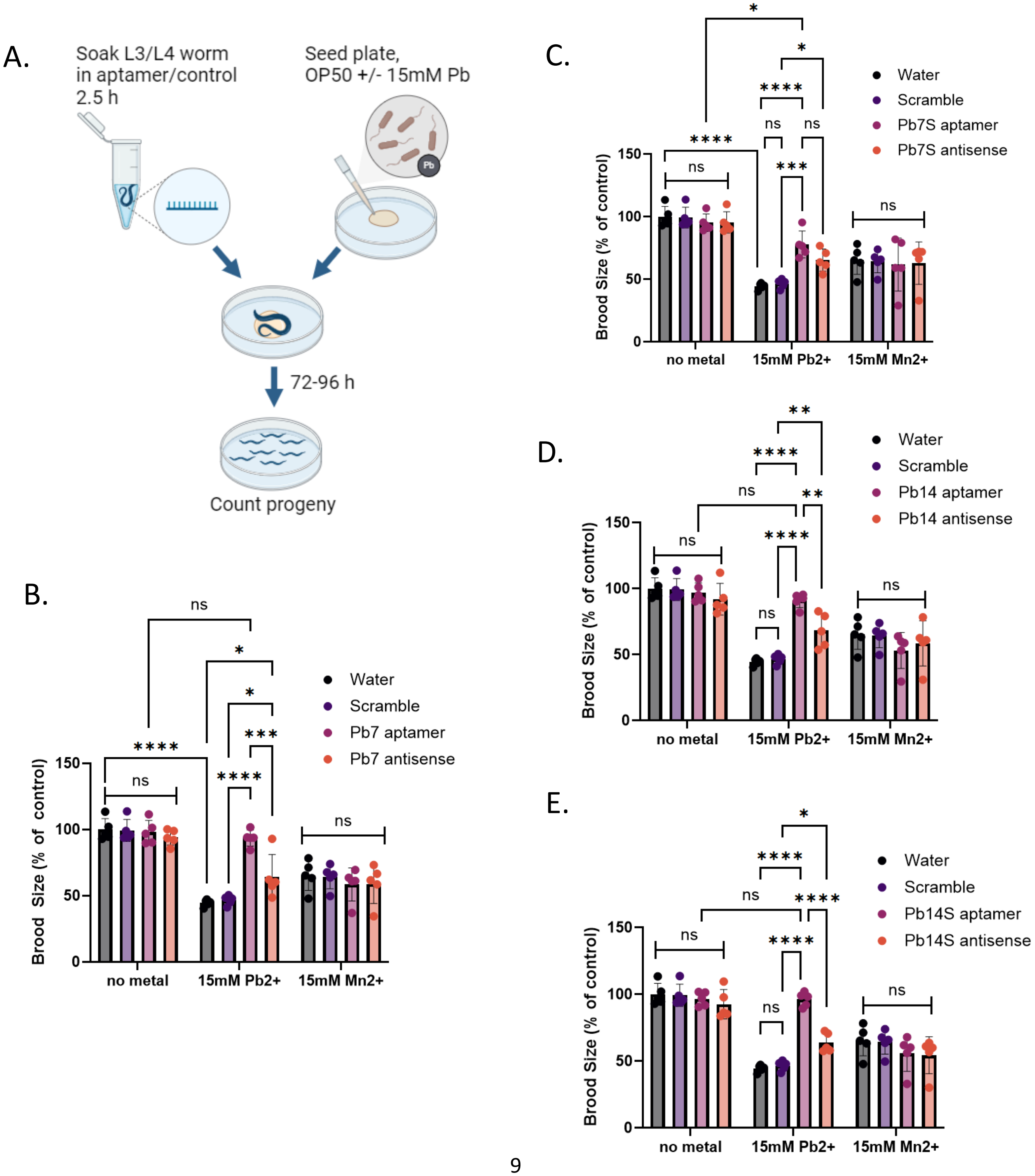

**Supplementary Figure S7. Cadmium induces a dose-dependent decrease in brood size.** Cadmium concentrations indicated were mixed with OP50 and plated to NGM plates, and brood sizes were established at 72 h as described. Error bars are +/- SEM, n=3.

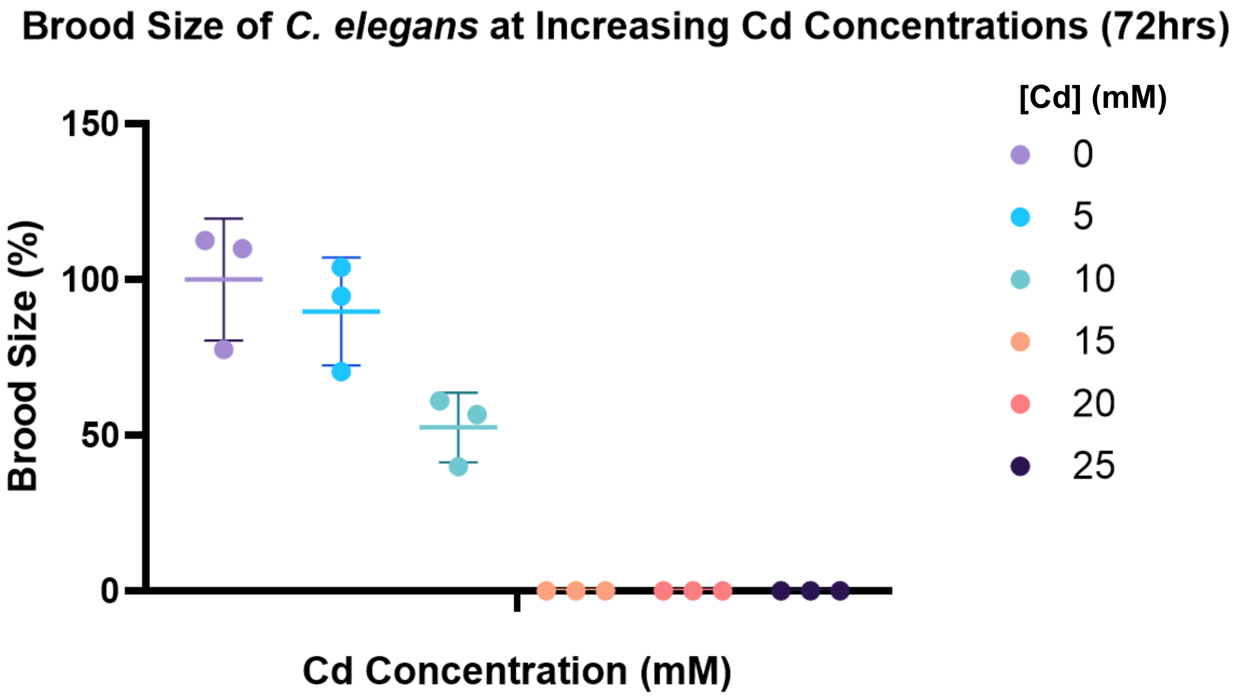

**Supplementary Figure S8. The Cd2-2 aptamer does not protect *C. elegans* from lead toxicity**. Error bars are +/- S.E.M., n=3. *p<0.05, **p<0.01.

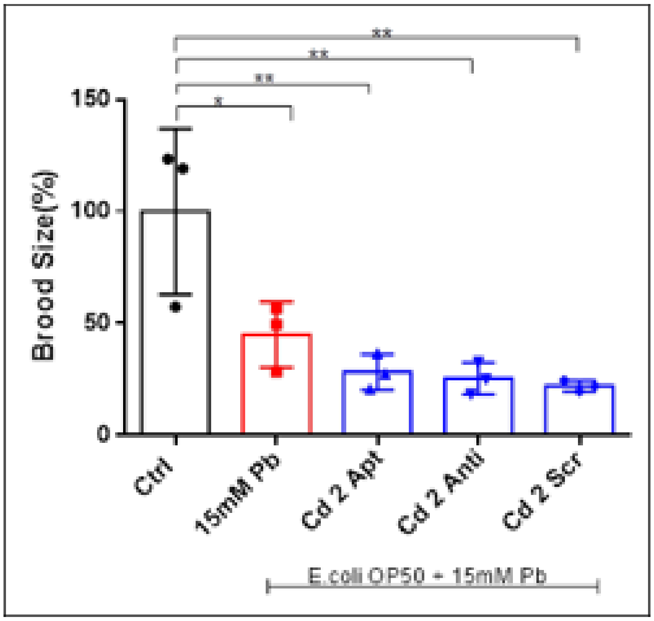
